# Assessment of bovine cortical bone fracture behavior using impact microindentation as a surrogate of fracture toughness

**DOI:** 10.1101/2023.08.07.552351

**Authors:** Babak Jahani, Rachana Vaidya, James M. Jin, Donald A. Aboytes, Kaitlyn S. Broz, Siva Khrotapalli, Bhanuteja Pujari, Walee M. Baig, Simon Y. Tang

## Abstract

The fracture behavior of bone is critically important for assessing its mechanical competence and ability to resist fractures. Fracture toughness, which quantifies a material’s resistance to crack propagation under controlled geometry, is regarded as the gold standard for evaluating a material’s resistance to fracture. However properly conducting this test requires access to calibrated mechanical load frames the destruction of the bone samples, making it impractical for obtaining clinical measurement of bone fracture. Impact microindentation offers a potential alternative by mimicking certain aspects of fracture toughness measurements, but its relationship with mechanistic fracture toughness remains unknown. In this study, we aimed to compare measurements of notched fracture toughness and impact microindentation in fresh and boiled bovine bone. Skeletally mature bovine bone specimens (n=48) were prepared, and half of them were boiled to denature the organic matrix, while the other half remained preserved in frozen conditions. Notched fracture toughness tests were conducted on all samples to determine Initiation toughness (K_IC_), and an impact microindentation test using the OsteoProbe was performed to obtain the Bone Material Strength index. Boiling the bone samples resulted increased the denatured collagen without affecting mineral density or porosity. The boiled bones also showed significant reduction in both K_IC_ (p < 0.0001) and the average Bone Material Strength index (p < 0.0001), leading to impaired resistance of bone to crack propagation. Remarkably, the average Bone Material Strength index exhibited a high correlation with K_IC_ (r = 0.86; p < 0.001). The ranked order difference analysis confirmed excellent agreement between the two measures. This study provides the first evidence that impact microindentation could serve as a surrogate measure for bone fracture behavior. The potential of impact microindentation to non-destructively assess bone fracture resistance could offer valuable insights into bone health without the need for elaborate testing equipment and sample destruction.

## Introduction

Catastrophic fractures are a significant public health problem with an alarming prevalence and substantial economic burden. Epidemiological data indicate that approximately 40% of women aged 50 and above will experience a fracture at the hip, vertebrae, or forearm in their remaining lifetime, with increased risk of complications and mortality^(1–3)^. Furthermore, it is projected that osteoporotic fractures in the United States alone will reach an annual incidence exceeding 3 million by 2025 and costs above 25.3 billion dollars per year^(4,5)^. This rise in fracture incidence can largely be attributed to progressive deterioration in bone strength and fracture resistance associated with aging and disease.

The fracture resistance of bone is influenced by multiple factors including bone geometry, density, and material properties of the bone matrix. Clinical imaging modalities such as X-ray and Dual Energy X-ray Absorptiometry (DXA) can robustly measure the geometry and metrics associated with mineral density^(6)^, however, directly assessing the mechanical properties of the bone matrix presents clinical challenges. Destructive mechanical testing methods, like bending, tension, compression, shear, indentation, and fracture toughness measurements, have been employed to characterize the mechanical competence of the bone matrix^(7–13)^. Each of these methods employ different mechanical principles and assumptions to analyze mechanical and material properties. To simulate a realistic failure of bone the fracture toughness method is considered the most definitive and realistic as it involves the controlled loading and expansion of a premade flaw.^(11,13)^. However, the destructive nature of these tests makes them impossible to implement in living organisms and humans.

In recent years, reference point indentation has emerged as a promising method for assessing the material behavior of bone tissue. Reference point indentation can fall under one of two categories: cyclic microindentation or impact microindentation. Cyclic microindentation, employs a reference probe that defines the reference position and then applies a trapezoid waveform with equal ramping, holding, and unramping that delivers successive indentations as the probe progressively penetrates the material. Cyclic microindentation provides a range of quantitative metrics that have demonstrated correlations with various bending behaviors observed in cortical bone tissue^(14,15)^. Conversely, impact microindentation (OsteoProbe) utilizes a reference force (rather than a reference position) to assess mechanical properties of cortical bone by measuring the normalized depth of indentation. The resultant impact microindentation measurements are associated with bending toughness^(16)^ and indentation hardness (e.g. Rockwell and Vickers Hardness tests)^(17)^ as well as compositional variations in bone tissue such as mineralization, collagen cross linking, and cortical porosity^(14,18,19)^. The application of reference point indentation technique to bone tissue enables the non-destructive assessments of mechanical behavior. Yet, the relationship between impact microindentation and fracture toughness measurements of the bone tissue has not been directly investigated. Thus, the objective of this study is to compare measurements of notched fracture toughness and impact microindentation in fresh and denatured bovine bone.

## Materials and methods

### Sample collection and preparation

Six bovine femurs were locally sourced from cows aged 12- to 18-months of age, the bones were wrapped in saline-soaked gauze and stored at −20°C prior to experimentation. From each femur, two beams were obtained per quadrant along the longitudinal axis (anterior, posterior, lateral, and medial) resulting in a total of forty-eight beams (Figure 1). The beams were cut and machined into rectangular parallelepipeds using a low-speed sectioning saw under constant water irrigation (UKAM, Industrial Superhard tools, Valencia CA, USA). The dimensions of the beams were as follows: Width (W) = 6 mm, Length (L) ∼50mm and thickness (B) = 3 mm.

**Figure 1:**
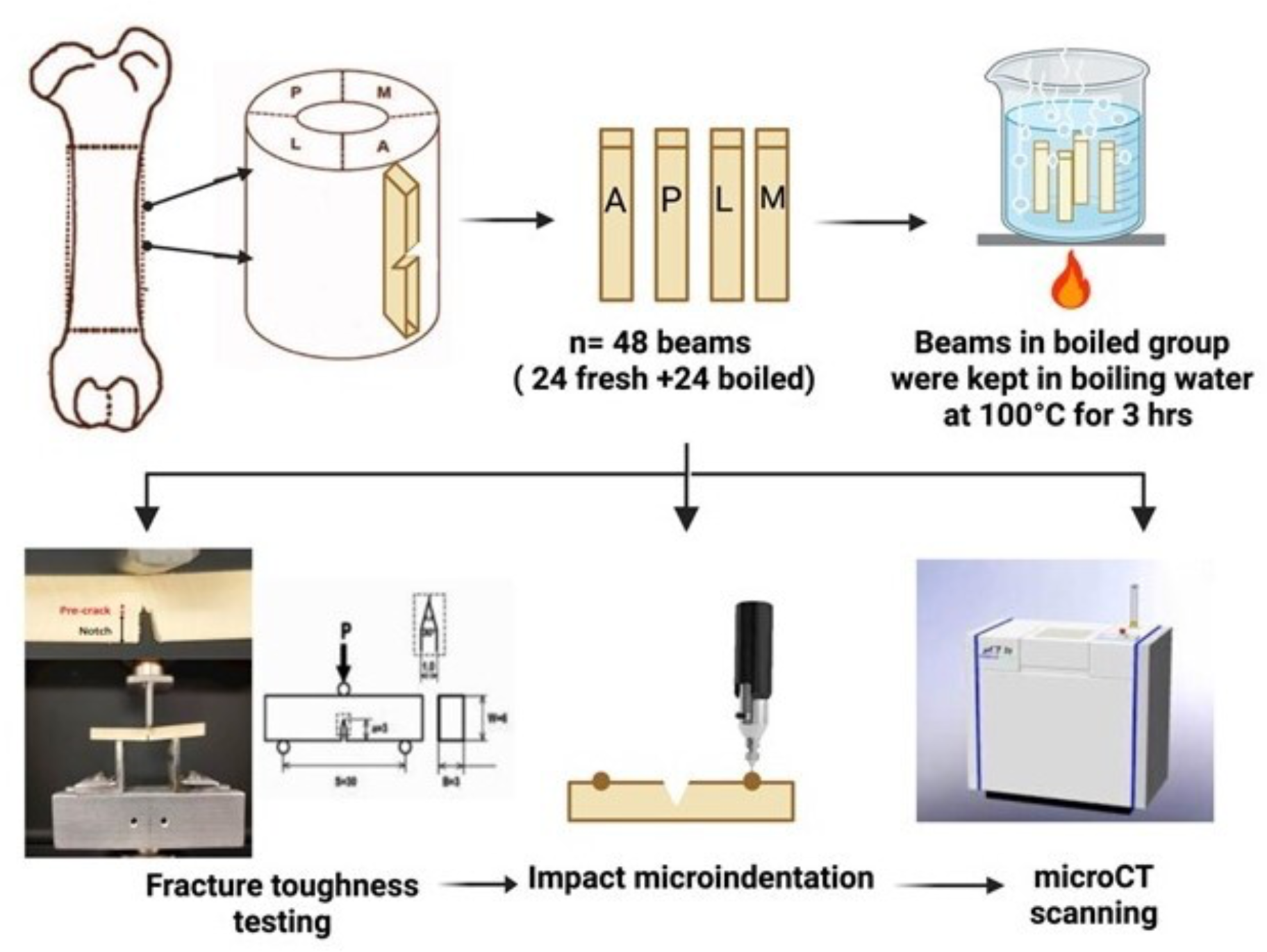
Overview of the experimental design. Forty-eight beams were extracted from six bovine femurs and paired by quadrant (Created with <u>BioRender.com</u>). All the beams were cut to the same dimensions of 50 mm x 6mm x 3mm. The boiled group underwent a 3-hour boiling while the control (fresh) group was stored in −20℃ until testing. Following the treatments, single edge notches were created on the periosteal side of each beam. Each notch was initiated using a diamond wafered blade to a depth of 1mm, then advanced to a pre-crack using a surgical blade with a polishing solution containing 0.3 μm diameter alumina suspensions. The notch width = 1 mm, point angle = 30°, and radius of the slit tip of 0.05 mm. The overall length of the notch and pre-crack was a=3 mm to maintain a/W=0.5. The beams subsequently underwent 3-point bending test with a preload of 10N and a loading rate of 0.2 mm/min until failure. Indentation measurements were performed using the OsteoProbe device, and the Bone Scores were reported as an average of 8 indentations. MicroCT scans were taken to evaluate the crack surfaces and the indentation sites.

For fracture toughness measurements, a single-edged V-notch and a pre-crack were introduced on the periosteal side of each beam. A diamond-wafered blade with a 30° taper was employed to create a 3 mm notch (the ‘a’ parameter) at the center of the beam. A pre-crack of 0.1 mm was introduced on each beam using a single-edge box cutter and Alumina suspension polish (0.3μm particle size; Allied High Tech, Compton, CA, USA). The dimensions of the notch and pre-crack combined approximated an aspect ratio of a/W=0.5.

Following machining and preparation, the beams were divided into two groups, with one bone of the pair assigned to be boiled and the other served as the control group. The boiled group was immersed in boiling water for a duration of 3 hours, while the control group remained stored at −20℃ until mechanical testing. Boiling the bone aimed to denature the organic matrix network components without impacting the mineral phase. All mechanical testing was conducted at standard room conditions (20℃, 50% humidity).

### Fracture toughness test

In accordance with ASTM E399 ^(20)^, the fracture toughness test was conducted in a three-point bending configuration (Span ‘S’= 30 mm, 1.5 mm support radius) using a Instron 5866 load frame (Instron Co, Norwood, MA). The samples were pre-loaded with 10N, similar to the 10N reference force of the impact indentation device, and then subjected to a loading rate of 0.2 mm/min until complete fracture, during which force-displacement data were collected. The critical stress intensity factor (K_IC_) of each beam was subsequently calculated using the following equation^(20)^:

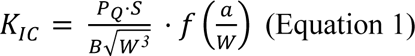

*P*_Q_: The maximum load of the linear region of load-displacement curve

S: Span of each beam under the load (30 mm)

B: Thickness of beam (3 mm)

W: Width of each beam (6 mm)

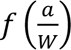 is the geometric correction term, given as follows:

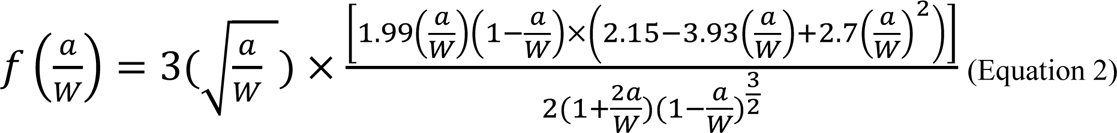

### Impact Microindentation test

Impact microindentation was performed on the periosteal side of each beam at least 20 mm away from the notch using OsteoProbe (Active Life Scientific, Santa Barbara, CA), a handheld device that is designed to measure the resistance of cortical bone to the indentation^(17)^. A reference load of 10N was manually applied by the operator using the measurement probe, and followed by application of a 30 N (total of 40N) load to the measurement surface at a rate of 120,000 N/s. The depth of penetration into the bone is recorded typically ranging from ∼150-400µm, and the outcome was reported as a singular scalar value, known as the Bone Material Strength index (BMSi)^(17,21,22)^. To calculate the BMSi value, eight indentations are made on the bone and the harmonic mean of the measured penetration distances into the bone is divided by the harmonic mean of ten penetration distances into a polymethylmethacrylate (PMMA) reference material and then multiplied by 100. PMMA measurements are made before and after each set of bone sample measurements.

### Measurement of Cortical Porosity and Tissue Mineral Density (TMD)

The 48 bovine beams were wrapped in PBS-soaked gauze and scanned using the µCT50 microCT scanner (Scanco Medical; Wangen-Brüttisellen, Switzerland) to evaluate tissue mineral density (TMD), cortical porosity, and resultant fracture surface. The entire half from each bone sample was scanned. The cortical bone analyses were obtained at a voxel size of 20 µm, while the fracture surface and indentation site visualization were obtained at 3 µm. The x-ray tube potential was 70 kVp with an intensity of 57 µA 500 projections and a 700 ms integration time.

TMD and cortical porosity values were derived from the cortical analysis module in the Scanco Evaluation software (v 1.2.30.0). Cortical porosity was calculated as the percentage of voxel bone volume divided by the total volume, with the final value subtracted from 100. 3D renderings of the bovine beams were generated by importing the CT image DICOMs into the 3D Slicer image analysis software (v 5.3.1, National Alliance for Medical Image Computing). The segment editor module was used to apply a threshold, determined by the distribution of air and bone attenuation constants, to segment the bone tissue from its surrounding, and then the fracture surfaces were rendered accordingly. The Scanco Visualizer software (v 1.2.13.0) was employed to visualize and qualitatively assess the crack surface, encompassing the region from the notch to the endosteal side of the beam^(23)^. The indentation regions were viewed from cross-sections that were created by aligning the virtual axis with the physical axis of the beam, and then successively sectioned until the largest indentation impression has been reached.

### Measurement of denatured collagen

The amount of denatured collagen in a subgroup of bone specimens (n=6 in each group) was quantified using previously described protocol^(24)^. Bone samples measuring 2 x 2 x 2 mm were collected from both the fresh and boiled groups and subjected to demineralization with Immunocal solution (Statlab, McKinney, TX). Each sample was then treated twice with extraction buffer containing 1 ml of 4M Guanidine hydrochloride in incubation buffer (0.1M Tris HCl, pH 7.3) containing protease inhibitors, iodoacetamide (1mM), and EDTA (1mM) for a duration of 48 hours at 4°C on a roller plate. This step ensured removal of proteoglycans and soluble collagen, and irreversibly denature any crosslinked collagen molecules that have suffered proteolysis or destruction due to excessive mechanical loading. Following the extraction, the bone samples were washed three times with incubation buffer for 3-6 hours at 4°C. The insoluble matrix, which contained the denatured collagen, was subjected to an overnight digestion at 37°C with incubation buffer containing 1 mg/ml of alpha chymotrypsin. The resulting supernatant containing the alpha chymotrypsin-solubilized collagen fragments was quantitatively collected and mixed at a 1:1 ratio with 6N HCl. Simultaneously, the remaining bone tissue was immersed in 6N HCl. Both the supernatant and the residual tissue were hydrolyzed at 110°C for a period of 20-24 hours. Following hydrolysis, the excess HCl was evaporated from the hydrolysates. The samples were then reconstituted with 500 μl of 0.1X PBS. Collagen content was quantified using the hydroxyproline assay, as previously described ^(25)^. Briefly, both the samples and hydroxyproline standards (stock: 200 μg/mL L-hydroxyproline) were incubated with chloramine-T solution at room temperature to facilitate the oxidation of hydroxyproline. Perchloric acid (3.15 M) was added to quench any residual chloramine-T, followed by an incubation period at room temperature. A p-dimethylaminobenzaldehyde solution was then added as a colorizing solution, and the mixture was incubated at 60 °C. All standards were allowed to cool at room temperature under complete darkness and the absorbance was measured at 560 nm using a synergy HTX microplate reader (BioTek, Winooski, VT). The amount of denatured collagen was expressed as the percentage of total amount of collagen.

### Statistical methods

The measured outcomes were plotted as histograms to examine distributions. Data remained paired and were compared between groups by Repeated Measures analysis of variance (ANOVA). Pearson’s correlations were run to identify the linear correlation between fracture toughness K_IC_ and Bone score (BMSi). Respective measurements were then ranked from high to low in order to perform a ranked order difference analyses^(26)^ to identify the systemic bias between Bone Score (BMSi) and fracture toughness (K_IC_). All analyses were conducted using PRISM 9 (GraphPad, San Diego, USA) or Excel (Microsoft, Redmond USA).

## RESULTS

### Boiling does not affect cortical porosity nor tissue mineral density, but it leads to a significant increase in the denatured collagen

The cortical bone porosity analyses revealed no significant difference in cortical porosity (p=0.57) between the boiled and fresh bones. The mean porosities for the fresh group and the boiled group were 0.20% (± 0.32%) and 0.27% (± 0.40%) respectively. Similarly, no significant differences were observed in tissue mineral density between the two groups (Figure 2B). The mean TMD values for the fresh and boiled group were 1005 (± 25.93) mg HA/cm^3^ and 1011(± 31.44) mg HA/cm^3^, respectively. Taken together, these results confirm that boiling did not substantially affect the microstructure or mineral composition of the bone. The

**Figure 2:**
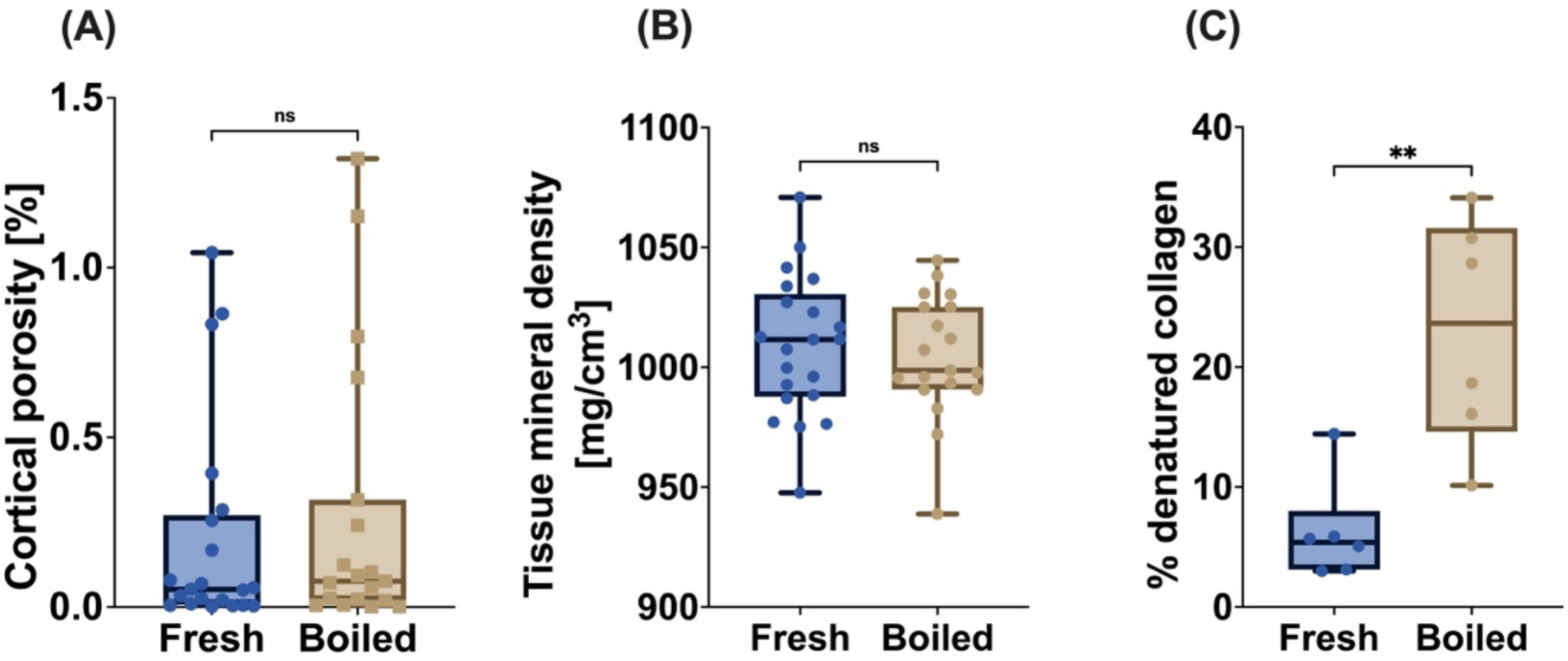
Boiling did not influence the (A) cortical porosity (p = 0.574) nor the (B) tissue mineral density (p = 0.462) of the bone beams. Samples were evaluated using microCT scanned at 20 μm spatial resolution. However, (C) boiling did increase the percentage of denatured collagen (p=0.005).

### Fracture toughness discerns the effects of boiling and quadrant-dependent variations of bone fracture behavior

The fracture toughness measurements of cortical bone in both fresh and boiled beams revealed a significant decrease in fracture toughness in boiled beams compared to fresh beams in all four quadrants of the bone (p<0.0001) (Figure 3A). We also observed an effect of quadrant on fracture toughness (p=0.002). Analysis of the fracture surfaces revealed distinct differences between the fresh and boiled samples (Figure 3B). Fresh beams exhibited a greater occurrence of crack deflection and toughening behavior, resulting in irregular and larger crack. (Figure 3B). These results confirm that boiling significantly deteriorates bone material properties and compromises the bone’s resistance to fracture.

**Figure 3:**
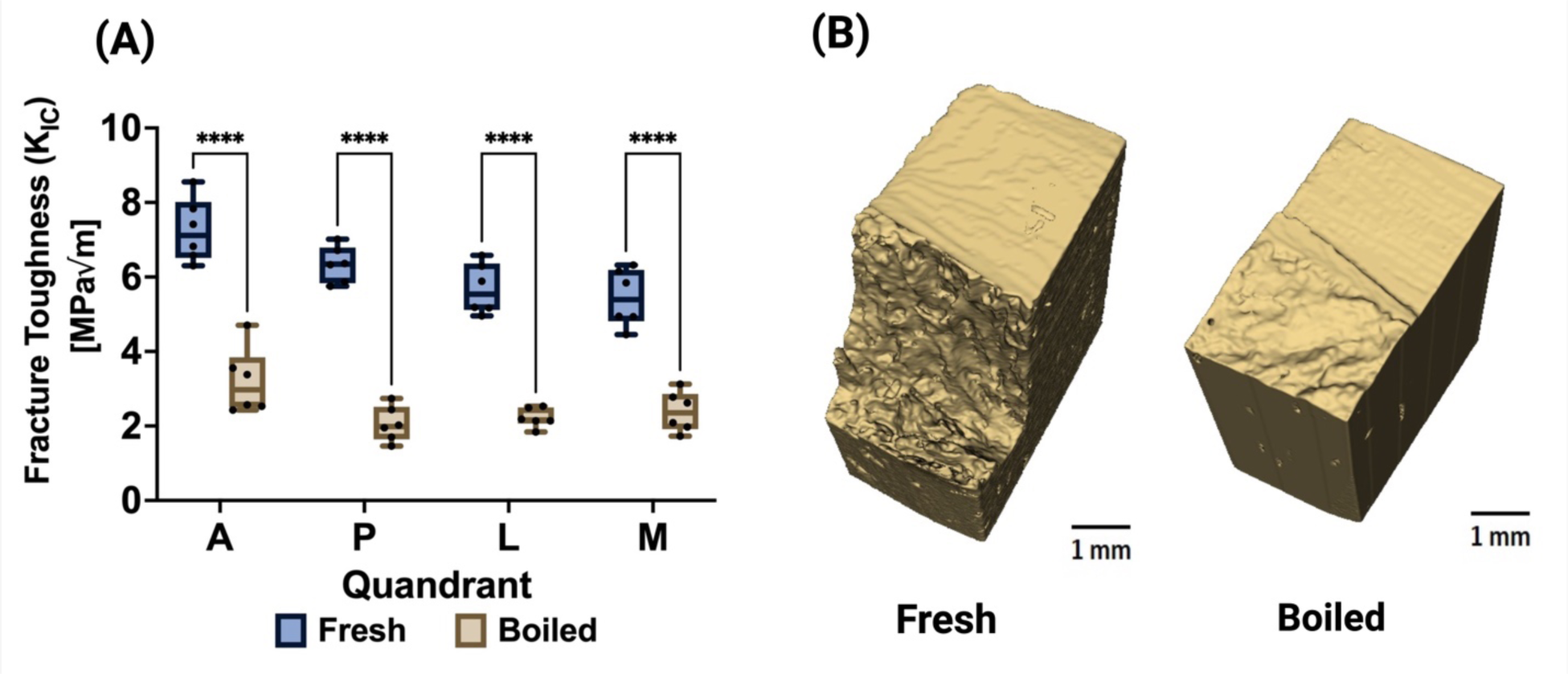
(A) Fracture toughness measurements show that fresh bovine beams had significantly higher fracture toughness (6.18 ± 0.81 MPa √m) than boiled beams in samples taken from all 4 quadrants (2.46 ± 0.507 MPa √m) (B) Inspection of the fracture surfaces from microCT revealed that the fresh bones had extensive crack deflection prior to failure, while the boiled samples showed little to no deflection of the crack. (**** p < 0.0001; *** p < 0.001; ** p < 0.01; * p < 0.05).

### Impact microindentation measurements are strongly correlated with initiation toughness

Impact microindentation was deployed to assess bone material properties in both fresh and boiled groups. The indentation measurements revealed a significantly lower average BMSi values in the boiled group, indicating detectable impairments in bone material caused by boiling (Figure 4A). No significant differences were observed between quadrants (p=0.160). High-resolution microCT illustrated that the reference load applied by the probe creates a observable impression similar to the stress concentrating pre-crack in the notched bone beams. Given the equal reference load applied on the bones of both groups, the boiled group incurred a larger and deeper impression (Figure 4B). Consistent with the BMSi readout, the subsequent indentation applied by the additional 30 N of force created a larger impression in the boiled group than in the fresh group. Importantly, a strong correlation was identified between the average BMSi and increasing fracture toughness (K_IC_) (Figure 5A). The ranked order difference analysis revealed that approximately 43 of the 48 measurements were within the Upper Acceptable Limits and Lower Acceptable Limits defined by the 95% confidence interval. Moreover, the bias converges towards zero where the fragile bones are ranked (Figure 5B).

**Figure 4:**
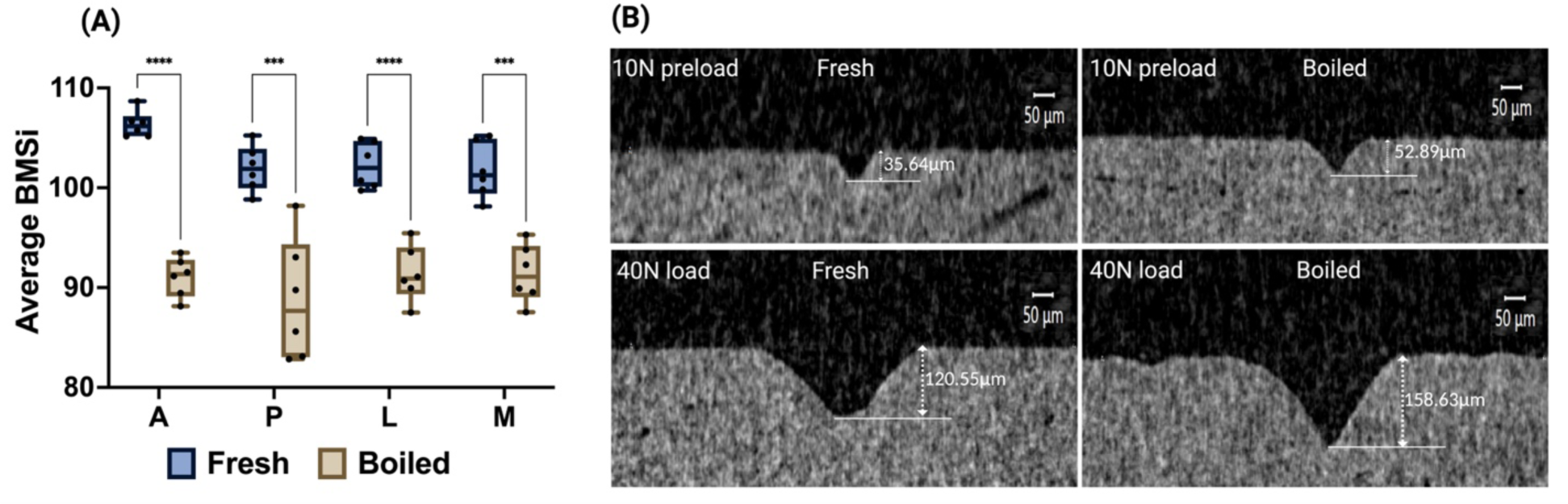
(A) Impact microindentation results showed that the average BMSi is significantly lower in the boiled group (90.6 ± 1.27 [unitless]) compared to the fresh group (103 ± 2.19 [unitless]) (p<0.0001) (B) MicroCT scans at 3μm resolution show that fresh beams show smaller indentation depths, 35.64 μm and 120.55 μm; compared to boiled beams, 52.89 μm and 158.63 μm, after preload of 10N, and full load of 40N, respectively.

**Figure 5.**
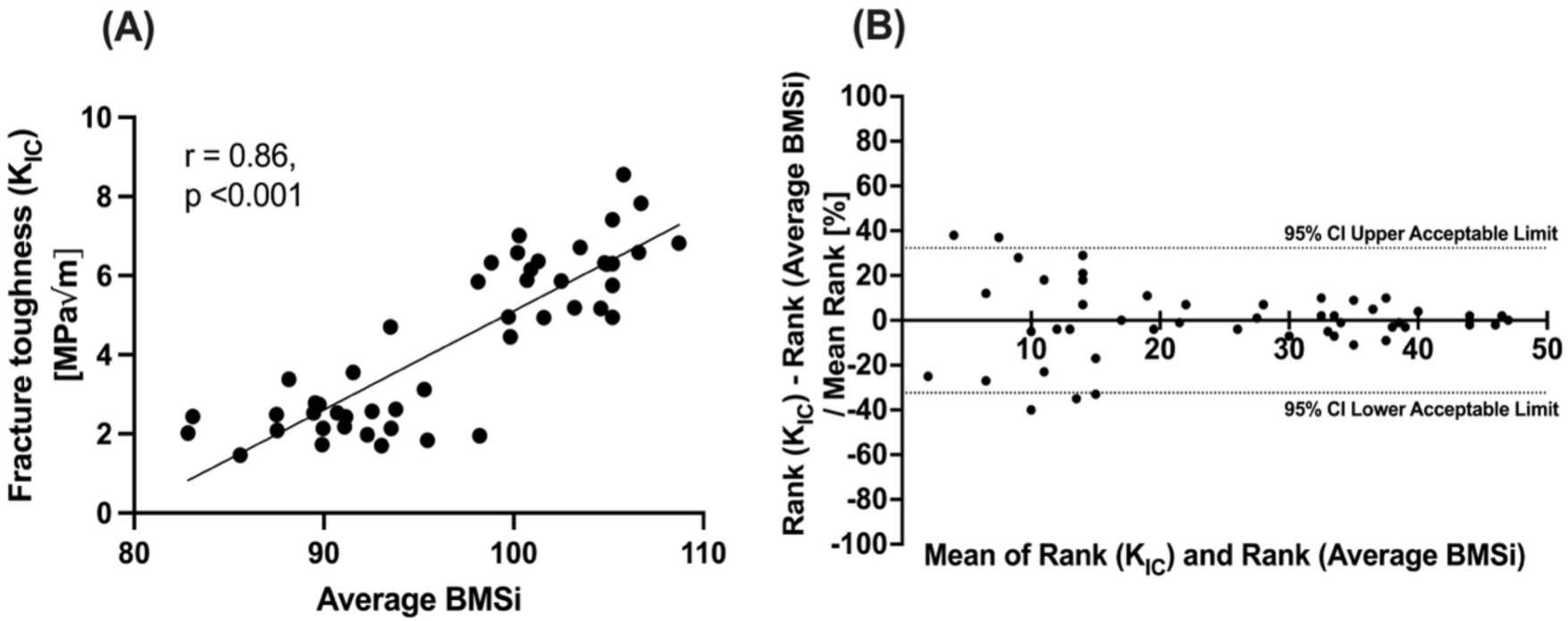
(A) The Bone Score (Average BMSi) measurements are strongly correlated to fracture toughness (K_IC_) (r=0.86, p<0.001). (B) The ranked difference analysis reveals approximately 90% of measurements lie well within the 95% CI Acceptable Limits. Moreover, the bias converges towards zero at the higher ranks where fragility occurs in these samples. This data suggests that Bone Score is an outstanding surrogate for fracture toughness K_IC_ in weakened bone.

## DISCUSSION

In this study, we investigated the fracture behavior of boiled and fresh bovine cortical bone using impact microindentation and fracture toughness testing. We subjected the bones to boiling, which denatures the protein tertiary structures, as a controlled method to deteriorate bone’s organic phase without affecting the mineral phase^(27)^. MicroCT analyses confirmed that boiling altered only the organic phase as seen by elevated percentage of denatured collagen in boiled samples, without exhibiting any discernible influence on the mineral phase, shown here as no differences in cortical porosity or tissue mineral density between groups. While the mineral phase showed no differences we noted a significant reduction in mechanical properties, specifically fracture toughness in the boiled samples. The loss of fracture toughness in the boiled beams was accompanied by distinctive morphologies of the crack surface (Figure 3B). The boiled samples exhibited smoother crack surfaces that signifies brittle fracture behavior and rapid crack propagation with minimal plastic deformation. In contrast, the fresh samples exhibited more tortuous crack surfaces that are typically associated with ductile fracture behavior, characterized robust deflection mechanisms^(27,28)^. The critical stress intensity factor (K_IC_) was used as the measure of fracture toughness and is based on linear-elastic assumptions for the bone material. This assumption minimizes the contribution from plastic deformation and thus does not account for crack-growth toughening^(29–31)^. Therefore, these results likely underestimate the true fracture toughness of bone. In contrast, impact microindentation measurements include elements of elastic and plastic behavior and may be a more comprehensive evaluation of the material’s damage behavior. To our knowledge, this is the first study to examine the relationship between fracture toughness and microindentation measurements using the OsteoProbe. It is worth noting that the notched fracture toughness is conducted at the millimeter scale, and therefore likely detected the contributions from microstructural variations known to occur across bone quadrants^(32)^. On the other hand, the micrometer-scale indentation measurements are at a smaller length-scale and therefore were not affected by these features^(7)^. The excellent correspondence between K_IC_ and BMSi suggests that measurements of the OsteoProbe provide insights on the fracture toughness of bone. The ranked order difference analysis is the gold-standard for comparing two analytical measurements with disparate units^(26)^, and this analysis revealed the bias for approximately 90% of the measurements were within the acceptable limits defined by the 95% confidence interval. Moreover, the magnitude of bias diminished in lower K_IC_ and BMSi (higher ranked) samples. While these limits will need to be refined with additional clinical data, the bias range here is considered good to excellent^(33)^.

Bone fractures occur though a complex interplay of material and biological factors. However, clinical assessments of fracture risk typically do not account for the material behavior of bone. The lack of clinical information of in vivo bone material quality highlights the importance of measuring bone fracture properties for fracture risk prediction. For examples, bone mineral density (BMD), measured by DXA, is currently considered the gold standard for determining fracture risk, yet, BMD has been shown to explain only one-third of an individual’s fracture risks in middle-aged and elderly women^(34,35)^. Since bone is a complex biocomposite, its fracture behavior is determined by a number of bone density-independent factors including degraded collagenous matrix, increased porosity, accumulation of microcracks^(14,30,36–40)^, among others. Combining BMD and material-level measures such as indentation improves the prediction of bone strength^(40)^. Impact microindentation methods have been successful in detecting changes in bone composition^(41)^, adaptations to bone therapeutics^(42–44)^, and pathological bone diseases^(19,41–43,45–56)^ with BMSi reported to be lower in the respective pathological populations. Our findings here suggest that these differences detected by indentation likely reflect the changes in bone fracture toughness in these patients.

It is also important to note that changes in BMSi do not always correlate with prevalent fractures^(57–60)^. Bone with impaired fracture resistance is more likely to fracture *ceteris paribus*, but whether a fracture event occurs can vary with individual loading history, exposure to traumatic loads, and current bone metabolism. Therefore, the ability to measure bone fracture resistance in a patient should be interpreted in context of their individual propensity and other risk factors. Nonetheless, *in vivo* fracture toughness of bone is undoubtedly a critical aspect of fragility especially in susceptible individuals.

There are a number of limitations that should be noted in this study. We used bovine cortical bone in our experiments, which may not perfectly emulate the properties of human bone. Although bovine bones are from skeletally mature animals, these animals are relatively young compared to elderly humans. Therefore, boiling to deteriorate bone’s organic phase produces dramatic changes and may not recapitulate the progressive biological changes that occur in bones due to aging and disease.

In summary, our findings demonstrate that impact microindentation can detect differences caused by heat denaturation, and these measurements closely correspond to the critical stress intensity factor measurement from a notched fracture toughness test. Although it is not possible to perform direct measures of fracture toughness in vivo, in vivo measurements of impact microindentation^(61)^ on bone could be considered a surrogate measure of bone fracture toughness.

## CONCLUSION

Our findings support that impact microindentation quantifies changes in bone fracture behavior. Bone material strength index was highly correlated with the critical stress intensity factor of fracture toughness measured in boiled and fresh bovine bone.

## ACKNOWLEDGEMENTS

This study was supported by the National Institutes of Health (P30 AR074992, R44 AG071034, R01 AR074441, R01 AR077678).

## Notes

### Competing Interest Statement

The authors have declared no competing interest.

### Summary of Updates

An additional analysis on measurement of denatured collagen is added to the manuscript. Changes were made in the methods section. Figure 2- addition of panel C, results section and discussion section to include data on denatured collagen. Author's list and affiliations updated.

